# CellCov: gene-body coverage profiling for single-cell RNA-seq

**DOI:** 10.64898/2026.01.30.702727

**Authors:** Shiyao Chen, Urša Zevnik, Christoph Ziegenhain

## Abstract

**Motivation:** Gene-body coverage bias differs across scRNA-seq protocols and can influence downstream analyses, yet coverage is often assessed using bulk-level summaries that obscure cell-to-cell variability.

**Results:** **CellCov** provides gene-body coverage profiling at single-cell resolution, enabling exploration of coverage heterogeneity across both cells and features. The accompanying workflow supports flexible grouping and robust aggregation of profiles by user-provided annotations, allowing principled comparison of coverage bias across sequencing protocols. We demonstrate its use on public datasets from several full-length scRNA-seq chemistries.

**Availability:** **CellCov** source code and documentation are available at https://github.com/ziegenhain-lab/CellCov

**Supplementary information:** Supplementary data are available at *Bioinformatics* online

## 1 Introduction

Single-cell RNA sequencing has become broadly accessible and is widely used in biomedical research. Recent technological advances have extended its use from simple gene-level counting of RNA tags (3’ or 5’ ends) to full coverage of transcripts (Ziegenhain *et al*. 2017; Mereu *et al*. 2020; Kumari *et al*. 2024; Molla Desta and Birhanu 2025). As such new methods continue to emerge, comprehensive benchmarking and appropriate QC are increasingly important to ensure validity of results.

A critical aspect of full-length RNA sequencing quality is gene body coverage, which describes how the sequencing reads are distributed from 5’ to 3’ end of the gene. Ideally, each position is sequenced with the same probability, leading to uniform coverage distribution (Li *et al*. 2015), but this is often not the case in practice. Common causes of 3’ bias include enzymatic fragmentation, incomplete reverse transcription, and mRNA degradation in combination with poly-A capture strategies. Conversely, 5’ bias can arise from fragmentation combined with 5′-end counting or internal oligo-dT priming (Chen, Ning and Shi 2019; Webber *et al*. 2025). Thus, gene-body coverage profiles not only depend on the protocol used but also reflect sample quality. It has been shown that skewed profiles are predictive of low-quality cells, and can be used for filtering them (Abugessaisa *et al*. 2022), highlighting the value of this metric in individual cells.

Nevertheless, tools commonly used for visualising coverage uniformity, e.g. RSeQC (Wang, Wang and Li 2012) and QoRTs (Hartley and Mullikin 2015), operate on bulk data. Although SkewC (Abugessaisa *et al*. 2022) is meant for analysis of single-cell data, it achieves this by splitting the input into per-cell BAM files, and running an RSeQC-like algorithm on each individually, which is computationally inefficient and at odds with the increasing scalability of single-cell approaches.

Moreover, comparing coverage profiles across genes or transcripts with different properties, such as GC content, expression level, or transcript length, is not possible with existing tools. Thus, detailed investigation of single-cell coverage profiles is currently not feasible. To overcome these limitations, we present CellCov, a lightweight tool that reports coverage with feature (gene or transcript) and cell resolution. This format enables flexible downstream analysis and facilitates comprehensive comparison at a single-cell level, as well as straightforward stratification across different feature properties.

## 2 Implementation

CellCov supports single-cell RNA sequencing data across a wide range of types (incl. short and long read data) and can also be applied to bulk RNA-seq datasets. The core component is a cell-barcode–aware base pileup engine implemented in Rust for computational efficiency, with a Python interface for preprocessing, postprocessing, and visualisation. CellCov requires alignments in BAM format, a list of cellular barcodes, and a GTF gene annotation file. For each annotated feature (gene or transcript), exonic coordinates are collected and mapped to gene-body percentiles (100 bins). The base counter processes one feature at a time and counts the number of aligned bases falling into each percentile bin for every barcode, producing a feature × cell coverage profile. Profiles are then stacked into a coverage tensor of shape (features × cells × 100) and L1-normalised per feature–cell pair to represent relative coverage along the gene body. CellCov additionally reports per-cell and per-feature summary statistics (e.g., coverage skewness) to support quality control and downstream stratification.

For user-defined cell and/or feature groups, CellCov can aggregate profiles into group-level gene-body coverage curves and optionally report variability bands to capture heterogeneity across cells or across features. The provided plotting script generates publication-ready figures and exports aggregated profiles for further analysis.

## 3 Application

To illustrate the utility of CellCov, we calculated coverage on public datasets generated from various scRNA-seq methods (Smart-seq3xpress (Hagemann-Jensen, Ziegenhain and Sandberg 2022), MAS-seq (Al’Khafaji *et al*. 2024), 10x Genomics 3’ Gene Expression v3, and R2C2 (Volden *et al*. 2018)) and explored the outputs with the provided plotting script.

CellCov outputs a coverage matrix with 100 percentile bins at single-cell single-feature resolution, which can be stratified by user-defined cell and/or feature annotations (e.g., cell type, gene length, GC content, expression) and aggregated with robust summaries to enable protocol comparisons. As an example, we binned genes by exonic length and jointly visualised all four datasets including cell-to-cell variability, showing clear chemistry-dependent length effects on coverage profiles (Fig. 1a). CellCov furthermore calculates a cellular metric of skewness, which is a directionless measure of how non-uniform coverage is distributed over gene bodies (Fig. 1b).

**Fig. 1.**
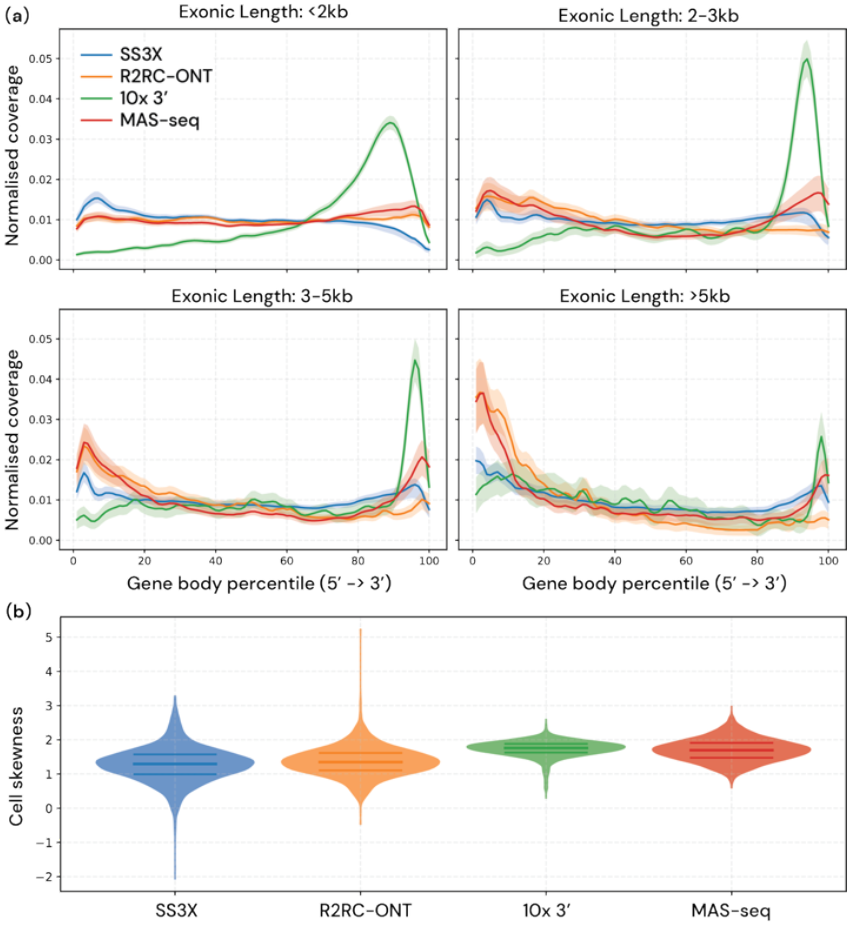
Gene-body coverage in single-cell RNA-seq protocols. (a) Normalised gene-body coverage (100 percentile bins from 5′ to 3′) is shown for genes grouped by exonic length (<2 kb, 2–3 kb, 3–5 kb, and >5 kb) across sequencing protocols (SS3X, R2RC-ONT, 10X, and MAS-Seq). Solid lines show the median coverage profile; shaded bands indicate cell-to-cell variability as the interquartile range (25th–75th percentiles). (b) Cellular coverage skewness is calculated and shown as violin plots.

## 4 Conclusion

CellCov is the first gene-body coverage calculation tool that produces perfeature and per-cell outputs from single BAM files. Its design allows computationally efficient base counting paired with custom filtering, grouping, and visualization directly on the main output, thus providing an unprecedented level of flexibility in method benchmarking and sample QC.

## Data Availability

Publicly available datasets used to generate Fig. 1 were downloaded from https://downloads.pacbcloud.com/public/dataset/MAS-Seq/ (Mas-seq), Smart-seq3xpress data is available at ArrayExpress under accession E-MTAB-11452, while 10x Genomics Illumina and R2C2 Oxford Nanopore Technologies data are available at SRA under BioProject accession PRJNA599962.

